# Improving robustness of 3D multi-shot EPI by structured low-rank reconstruction of segmented CAIPI sampling for fMRI at 7T

**DOI:** 10.1101/2021.08.19.457024

**Authors:** Xi Chen, Wenchuan Wu, Mark Chiew

## Abstract

Three-dimensional (3D) encoding methods are increasingly being explored as alternatives to multi-slice two-dimensional (2D) acquisitions in fMRI, particularly in cases where high isotropic resolution is needed. 3D multi-shot EPI is the most popular 3D fMRI acquisition method, but is susceptible to physiological fluctuations which can induce inter-shot phase variations, and thus reducing the achievable tSNR, negating some of the benefit of 3D encoding. This issue can be particularly problematic at ultra-high fields like 7T, which have more severe off-resonance effects. In this work, we aim to improve the temporal stability of 3D multi-shot EPI at 7T by improving its robustness to inter-shot phase variations. We presented a 3D segmented CAIPI sampling trajectory (“seg-CAIPI”) and an improved reconstruction method based on Hankel structured low-rank matrix recovery. Simulation and in-vivo results demonstrate that the combination of the seg-CAIPI sampling scheme and the proposed structured low-rank reconstruction is a promising way to effectively reduce the unwanted temporal variance induced by inter-shot physiological fluctuations, and thus improve the robustness of 3D multi-shot EPI for fMRI.

**Highlights:**

- A 3D multi-shot EPI sampling trajectory using interleaved ordering along *k_z_* and a CAIPI blipping pattern improves robustness to inter-shot phase variations
- Reconstruction based on Hankel structured low-rank matrix completion can significantly improve the temporal stability of 3D multi-shot acquisitions at 7T
- 1.5mm resolution brain fMRI data show that ~60% improvement in mean tSNR can be obtained using the proposed method compared to the conventional method
- 1.8mm resolution brain fMRI data demonstrate that the proposed method allows for 4-fold acceleration without loss of tSNR compared to conventional 3D EPI
- Preliminary brainstem fMRI data show that ~40% improvement in mean tSNR can be obtained using the proposed sampling and reconstruction

## 1. Introduction

Three-dimensional (3D) encoding methods are increasingly being explored as alternatives to twodimensional (2D) multi-slice acquisitions in functional MRI (fMRI). Compared to conventional 2D multi-slice imaging, 3D imaging is well known to provide SNR benefits as the whole volume rather than a thin slice is excited repeatedly for every shot. Also, 3D imaging is more favourable for reducing the total acquisition time for each volume (TR_volume_) by enabling flexible parallel imaging and partial Fourier acceleration along two phase encoding directions (Bilgic et al., 2019; Poser et al., 2010). In addition, 3D imaging can achieve high isotropic resolution without being limited by RF slice profiles along the partition/slice encoding direction. All of these advantages contribute to the prevalence of 3D encoding in high isotropic resolution fMRI, which has received strong interests for applications such as layer-specific fMRI (Lawrence et al., 2019). In particular, 3D multi-shot EPI, which samples a 2D *k_x_-k_y_* plane for each shot is one of the most conventional choices for 3D fMRI.

However, compared to 2D single-shot EPI, the main disadvantage of 3D multi-shot EPI imaging is that it suffers from increased vulnerability to shot-to-shot inconsistencies due to physiological fluctuations. The tSNR of a time series, defined as its temporal mean divided by temporal standard deviation, reflects the temporal stability of a given time-course, and relates to the sensitivity of the fMRI measurement. Ideally, tSNR is the same as the SNR of each volume, in the absence of physiological noise or scanner fluctuations. In practice, physiological noise scales with signal intensity and field strength, which creates an asymptotic limit on the achievable tSNR (Krüger and Glover, 2001; Triantafyllou et al., 2005). Inter-shot k-space inconsistencies can result in temporally varying ghost artifacts and thus a reduction in tSNR (van der Zwaag et al., 2012). Hence, the SNR benefits enabled by 3D encoding are not fully realized, and 3D multi-shot EPI might only be able to offer higher tSNR than 2D single shot EPI in low SNR, thermal noise dominated regimes (Jorge et al., 2013; Lutti et al., 2013).

These physiological fluctuations can manifest themselves as rather localized fluctuations of blood and cerebrospinal fluid related to cardiac pulsation, and spatially varying phase modulations resulting from the B0 fluctuations mainly caused by the movement of the chest during respiration (Tijssen et al., 2011; Zahneisen et al., 2014), the latter which is particularly troublesome for multi-shot imaging. Also, as the off-resonance effects scale with field strength (Triantafyllou et al., 2006), the physiologically induced inter-shot phase variations can be more detrimental at ultra-high fields like 7T, which plays an important role in high resolution fMRI due to its ability to boost SNR in small voxel regimes.

Previous work has shown that tSNR depends on the number of shots used to form a volume in a conventional 3D multi-shot EPI acquisition for fMRI, and that a larger number of shots is usually associated with a lower tSNR at the same SNR level (van der Zwaag et al., 2012). However, although the interaction between sampling trajectory and noise amplification (characterized by g-factors) has been well studied (Breuer et al., 2006; Narsude et al., 2016), and some work has previously explored the interaction between the ordering of k-space acquisition and physiological dynamics (Polimeni et al., 2016; Tijssen et al., 2011), the interaction between sampling trajectory and tSNR in the presence of non-trivial physiological noise contributions has not yet been fully explored.

Furthermore, while various methods have been proposed to correct physiological noise for fMRI, most of them are post-processing methods, either “model-based”(Agrawal et al., 2020; Glover et al., 2000; Hutton et al., 2011; Jorge et al., 2013; Kasper et al., 2017; Lutti et al., 2013) or “data-driven”(Jorge et al., 2013; Kasper et al., 2017; Salimi-Khorshidi et al., 2014), that work on reconstructed image time series. Unlike these post-processing methods, reconstruction methods which take into account the characteristics of multi-shot acquisitions generally rely on navigators to estimate the shot-to-shot phase variations (Barry et al., 2008; Barry and Menon, 2005). However, navigator techniques estimate dynamic phase information at the cost of prolonged acquisition time, and it can be particularly challenging to acquire 3D navigators. In the context of diffusion imaging, motion induced inter-shot phase variations due to the use of strong diffusion encoding gradients presents a similar problem for multi-shot acquisitions, and a variety of methods have been proposed to deal with this issue. These methods in general are also based on phase estimates (Chen et al., 2013). Recently, the MUSSELS (Mani et al., 2017) image reconstruction approach, which is based on Hankel structured low-rank matrix completion (Haldar, 2014; Jin et al., 2016; Shin et al., 2014), demonstrated the ability to account for inter-shot phase variations without knowing them explicitly, and has shown superior performance compared to explicit phase estimation methods.

This work aims to improve the temporal stability of 3D multi-shot EPI for fMRI at 7T, by reducing its vulnerability to physiologically induced inter-shot phase variations. Firstly, we explore the impact of sampling trajectories on the vulnerability to inter-shot phase variations and present a segmented 3D CAIPI sampling trajectory “seg-CAIPI”, which has the potential to improve tSNR in certain acquisition regimes. We also propose a reconstruction method based on Hankel structured low-rank matrix recovery, using an adaptation of the MUSSELS approach, to reduce the physiologically induced k-space inconsistency in 3D multi-shot EPI. Both simulation and in-vivo experiments demonstrate that the combination of the seg-CAIPI sampling trajectory and the structured low-rank reconstruction method provides a robust way to improve the tSNR of 3D multi-shot EPI imaging for fMRI.

## 2. Methods

### 2.1 seg-CAIPI sampling trajectory

The conventional 3D multi-shot EPI trajectory for which each shot samples a single partition and different partitions are sampled sequentially, is shown in Fig 1a, and it is referred to as “standard” trajectory in this work. We introduce a 3D multi-shot EPI trajectory termed “seg-CAIPI”, which uses interleaved sampling along *k_z_*, combined with *k_z_*-blipped CAIPI pattern (Breuer et al., 2005; Breuer et al., 2006; Setsompop et al., 2012), as shown in Fig.1b. In Fig. 1, each sample represents a readout line and all the samples connected by a dashed line correspond to a single shot. The index numbers above the trajectory diagram indicate the order in which different shots are sampled. In whole brain high resolution acquisitions, a large number of shots are required to sample an entire 3D k-space. In order to reduce the shot-dimensionality, we introduce a concept of “aggregate segment”, a binning of consecutive shots which are deliberately chosen to have an approximately uniform under-sampling pattern. For brevity, we refer to these binned consecutive shots as “segments” hereafter. This simplifies the problem by assuming the intra-segment inconsistencies are negligible, as respiratory effects (~3-5s) are expected to be temporally coherent on the timescale of each segment (~0.5s). As mentioned before, the inter-segment phase inconsistencies lead to temporally varying ghost artifacts, which contribute to unwanted temporal signal fluctuations.

**Figure 1.**
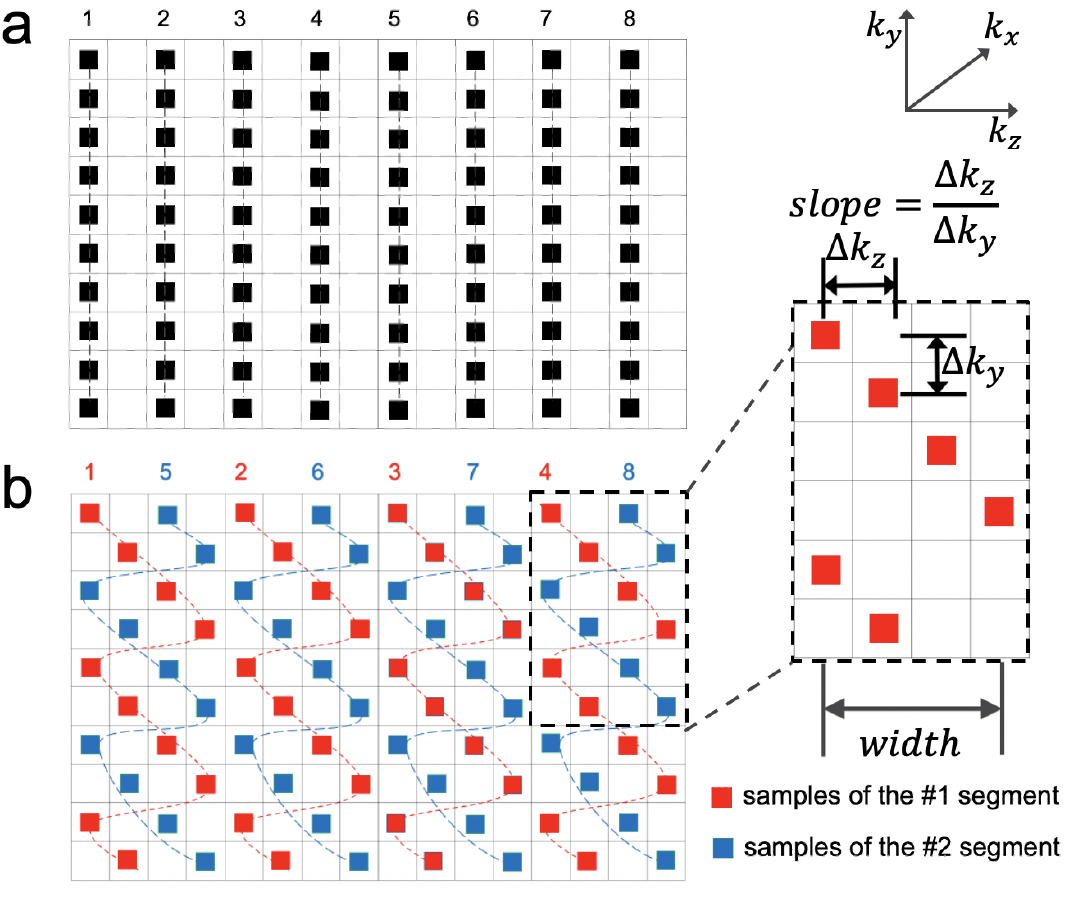
Schematic 3D multi-shot EPI trajectory shown in k_y_-k_z_ plane. Each sample represents a readout line and all the samples connected by a dashed line correspond to a single shot. Different shots are sampled in the order indicated by the index number above the trajectory diagrams. (a) A commonly used trajectory which we refer to as standard trajectory. All the shots are sampled sequentially along k_z_ direction. (b) The proposed seg-CAIPI trajectory. Different shots are sampled in an interleaved manner and k_z_ CAIPI blips are used for each shot. In this example, the first four shots are binned into a single segment (red samples) and the next four shots are binned into another segment (blue samples).

The seg-CAIPI sampling scheme is characterized by two parameters. The “width” parameter describes the offset between two consecutive shots within a segment which is chosen according to the number of shots per segment, and the “slope” parameter characterises the CAIPI pattern which indicates the ratio between the sampling step size along *k_z_* and *k_y_*. As the k-space raster loops back through previously sampled k-space regions across shots, a sampling offset along *k_z_* chosen to approximate a golden ratio step (~0.618 × *width*) was used to facilitate sampling uniformity. Note for both standard and seg-CAIPI trajectories we also use a Δ=1 blip in *k_y_* direction across consecutive shots when acceleration along PE dimension is employed, to stagger the PE lines for more distributed *k_y_* encoding. The standard trajectory is equivalent to the seg-CAIPI trajectory without *k_z_* blip and with only one segment.

### 2.2 Data Acquisition

All in vivo data were collected on a Siemens Magnetom 7T scanner (Siemens Healthineers, Erlangen, Germany) equipped with a 32-channel head-only receive coil with single-channel transmit (Nova Medical, Wilmington, MA, USA). Under-sampling was applied along two dimensions with the total acceleration factor *R* = *R_PE_* × *R*_3*D*_, where *R_PE_* denotes the acceleration along phase encoding direction for each shot and *R*_3*D*_ denotes the reduction on the number of shots acquired per volume. For seg-CAIPI under-sampling, *R*_3*D*_ is achieved by keeping the number of shots per segment fixed while reducing the number of segments acquired, i.e., for a 2x acceleration we only acquire the first 50% of shots needed to fully sample k-space. The specific trajectory parameters used in a given seg-CAIPI readout are specified as “seg-CAIPI(width, slope)”.

### 2.3 Low-rank constrained reconstruction

Unlike conventional reconstruction methods which combine all the acquired shots in k-space directly regardless of the shot-to-shot inconsistencies, we proposed a reconstruction method which aims to jointly reconstruct individual images for each segment. Images from different segments are then sum-of-squares combined to avoid phase cancellation effects. Similar to the Hankel structured low-rank matrix recovery methods MUSSELS (Mani et al., 2017), ALOHA (Jin et al., 2016; Lee et al., 2016a; Lee et al., 2016b), LORAKS (Haldar, 2014; Haldar and Zhuo, 2016), and SAKE (Shin et al., 2014) the missing k-space data are jointly recovered by exploiting the linear dependency among segments. To enable this, we assume that the images for each segment have the same magnitude, and differ only by a physiologically induced phase modulation. This redundancy in multi-segment k-space can be expressed as the low rank property of its block Hankel structured matrix representation. The reader is referred to reference (Mani et al., 2017) for a more detailed description of this property, although the reconstruction model also leverages all other linear inter-dependencies (such as those arising from limited image support) simultaneously through the structured low-rank matrix formulation (Haldar and Setsompop, 2020). Figure 2 shows one way to construct a block Hankel structured matrix from the multi-segment k-space. Then, image reconstruction is formulated as a low-rank constrained parallel imaging optimisation problem:

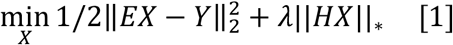

where *X* is the coil-combined multi-segment k-space to be reconstructed and *Y* is the measured multichannel multi-segment data. The forward model *E* performs the composition of *M* · *F* · *S* · *F*^−1^, where *F* and *F*^−1^ are the Fourier and inverse Fourier transform respectively. *S* denotes the multi-coil sensitivity encoding operator, and *M* selects the sampled k-space locations for each segment. The operator *H* applied to *X* generates the block-Hankel representation of *X*. ∥*HX*∥_*_ denotes the nuclear norm of *HX*, which is the convex approximation of *rank*(*HX*). *λ* is the regularisation parameter which weights the low-rank constraint. Eq.1 is solved using the Alternating Direction Method of Multipliers (ADMM) (Boyd, 2010) algorithm, which reformulates it as a constrained problem of the form:

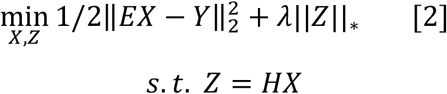

**Figure 2:**
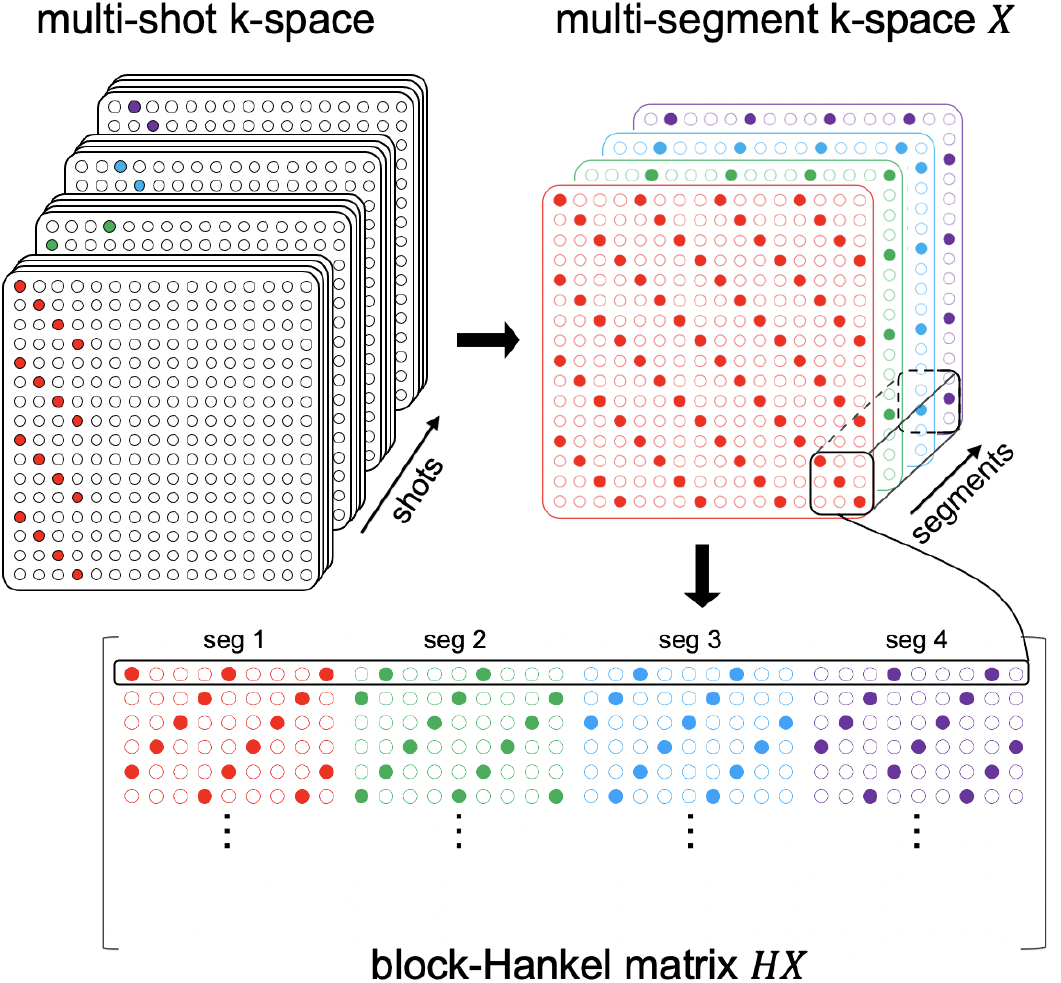
Illustration of the multisegment k-space data generation and the block-Hankel matrix construction from multi-segment k-space data. A small number of consecutive shots are binned together as a segment. X denotes the multi-segment k-space data and HX gives the block-Hankel matrix representation of it. A small patch of X selected by a sliding window is vectorized to generate a row of the block-Hankel matrix.

All reconstructions were implemented in MATLAB R2016a (MathWorks, Inc.). Coil sensitivity maps were calculated using ESPIRiT implemented in the BART toolbox (Uecker et al., 2014). The kernel size for Hankel transform was chosen empirically to be 6 × 6, and we used a fixed number of 10 iterations for the ADMM optimization. Reconstruction hyperparameters were also chosen empirically to maximize the resulting tSNR. Since image artifacts arising from shot-to-shot inconsistencies do not occur in the readout direction, an inverse Fourier transform was performed along the fully sampled readout direction, followed by separate reconstruction of each 2D *k_y_-k_z_*slice, which reduced the computational burden. Data acquired with the standard trajectory are not compatible with the proposed reconstruction due to the clustered nature of sequentially acquired shots, and were reconstructed by conventional methods, either a direct inverse Fourier Transform on fully sampled data, or SENSE reconstruction (Pruessmann et al., 1999) on the under-sampled data.

We evaluated the performance of the proposed sampling trajectory and reconstruction method by comparing three combinations of sampling and reconstruction:

1. standard: standard trajectory data with conventional reconstruction;
2. seg-CAIPI + conventional: seg-CAIPI trajectory data with conventional reconstruction;
3. seg-CAIPI + proposed: seg-CAIPI trajectory data with the proposed structured low-rank reconstruction.

While the “standard” data serves as a baseline reference for the impact of the sampling trajectory, the comparison between “seg-CAIPI + conventional” and “seg-CAIPI + proposed” demonstrates the impact of the structured low-rank reconstruction on identically sampled data.

### 2.4 Code and Data Availability Statement

All reconstruction code is available at https://github.com/XChen-p/Muitishot-EPI.

### 2.5 Simulation experiments - Impact of thermal noise and acceleration factor

Multi-shot k-space datasets consisting of a single *k_y_-k_z_* plane were synthesized by modulating a ground truth image with a series of physiologically plausible phase variation maps, which were measured from a 2D EPI time series acquired in sagittal orientation at 7T with 1.5mm isotropic resolution and TE/TR=20/40ms. The measured phase variation maps were then fit to a second order spherical harmonic basis set for denoising. The fully sampled k-space data for each shot was generated by resampling the 2D k-space time-series for both the standard trajectory and the seg-CAIPI(8,1) trajectory. To simulate low/medium/high thermal noise to physiological noise ratios, different levels of complex, zero-mean Gaussian noise were then added to the above k-space data respectively. Three physiological noise free datasets with the same amount of Gaussian noise were also generated as the reference datasets, whose mean SNRs are around 97/65/43 and indicate the tSNR upper bounds of the physiologically corrupted datasets. The reference datasets sampled by the standard and seg-CAIPI(8,1) trajectories were both generated. At the medium thermal noise level, multi-shot k-space datasets with varying *R_PE_* factors were also simulated.

### 2.6 In-vivo experiments

Three healthy volunteers were scanned at rest in accordance with local ethics to assess the tSNR performance of the proposed methodology. We investigated the impact of spatial resolution (1.2/1.5/1.8 mm isotropic resolution), trajectory parameters “width”and “slope”, and the acceleration parameter *R*_3*D*_, on the performance of the proposed method in terms of its ability to improve tSNR. An additional subject was scanned in a block-design visual checkerboard experiment to assess the impact on BOLD activation maps, using a 30/30s off/on block design. The functional datasets were processed by FEAT tool as part of FSL (the FMRIB Software Library) (Jenkinson et al., 2012; Woolrich et al., 2001), and minimal data pre-processing was used which includes high pass filtering and motion correction by MCFLIRT (Jenkinson et al., 2002).

A summary of the scanning protocols for all subjects are shown in Table 1. All pairs of standard and seg-CAIPI trajectory data used for comparison were acquired with matched parameters.

**Table 1.**
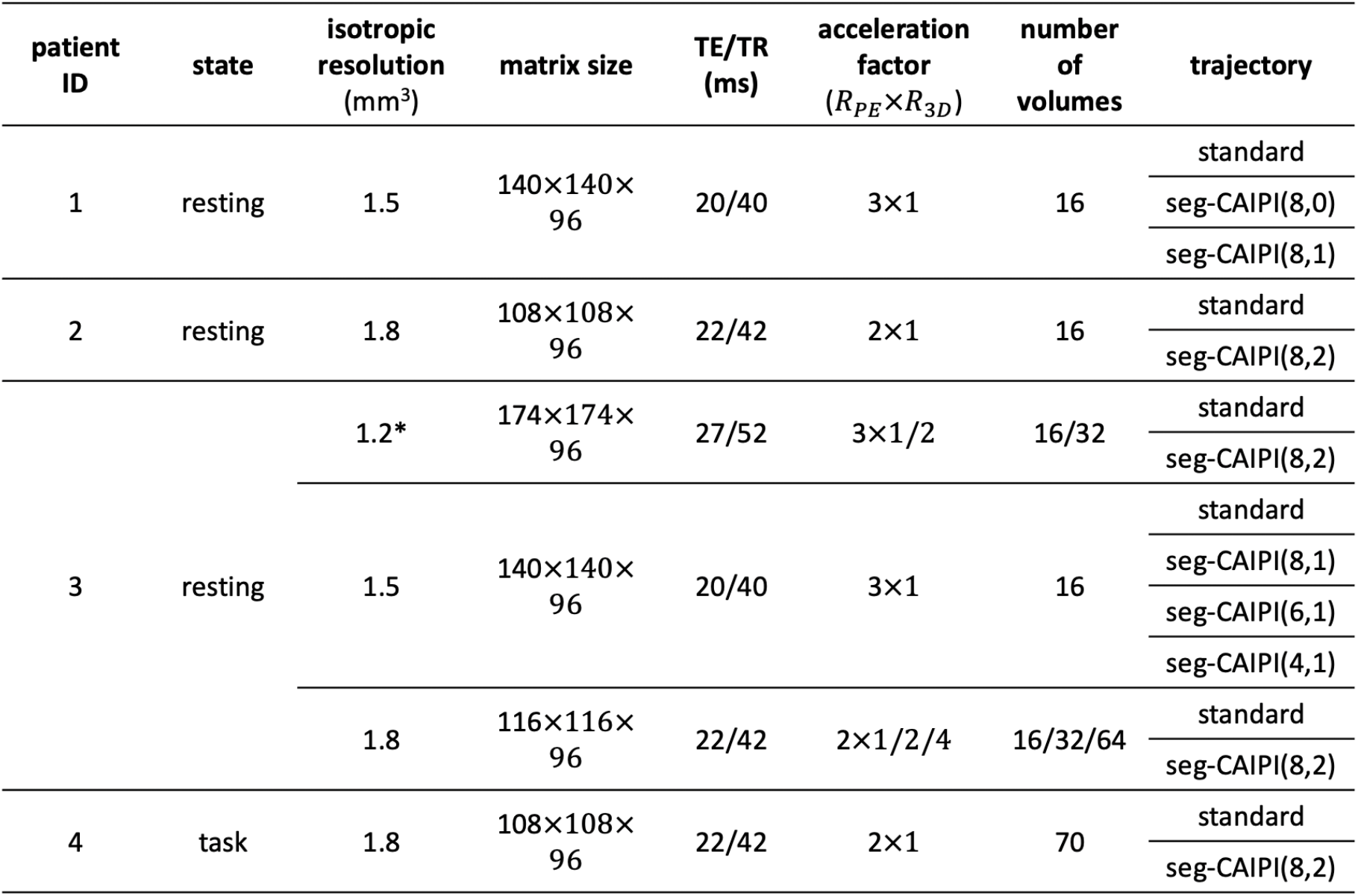
The scanning protocols of the in-vivo experiments. Cerebrum and brainstem imaging are both performed respectively on subject 3 with the protocol marked by *.

## 3. Results

Figure 3 shows the results of the simulation experiments. Fig 3a compares the mean tSNR for different methods at low/medium/high thermal to physiological noise ratios. With the conventional reconstruction, the seg-CAIPI data achieves higher tSNR than the standard acquisition at all three thermal noise levels, despite identical overall k-space coverage. With the proposed structured low-rank reconstruction on the seg-CAIPI data, further gains in tSNR are observed, achieving tSNR levels approaching the thermal noise only reference. As expected, when thermal noise increases, the relative tSNR differences between methods are reduced. Fig 3b shows the impact of acceleration factor *R_PE_* on the performance of different methods at medium thermal noise levels. The tSNR advantage of the seg-CAIPI trajectory under conventional reconstruction diminishes with *R_PE_*, performing on par with the standard approach at *R_PE_* = 3. In contrast, the proposed reconstruction on the seg-CAIPI data retains a significant tSNR improvement in all cases. Note the two thermal noise only reference datasets with the standard and seg-CAIPI trajectories show almost the same tSNR in both *R_PE_* = 2/3 under-sample acquisitions, which indicates the tSNR difference between these two sampling trajectories is mostly contributed by their varying robustness to physiological noise. Fig 3c shows images comparing the conventional (left) and proposed (right) reconstruction of the seg-CAIPI data at medium thermal noise level, by showing the maps of tSNR, temporal mean, temporal standard deviations, and temporal correlations with the reference data. The proposed reconstruction achieves an 18% improvement in mean tSNR compared to the conventional reconstruction, mainly by reducing the temporal standard deviation, as evidenced by the similar temporal mean images. In areas where temporal standard deviation is reduced by the proposed reconstruction, the correlation with the reference dataset time course is also improved.

**Figure 3.**
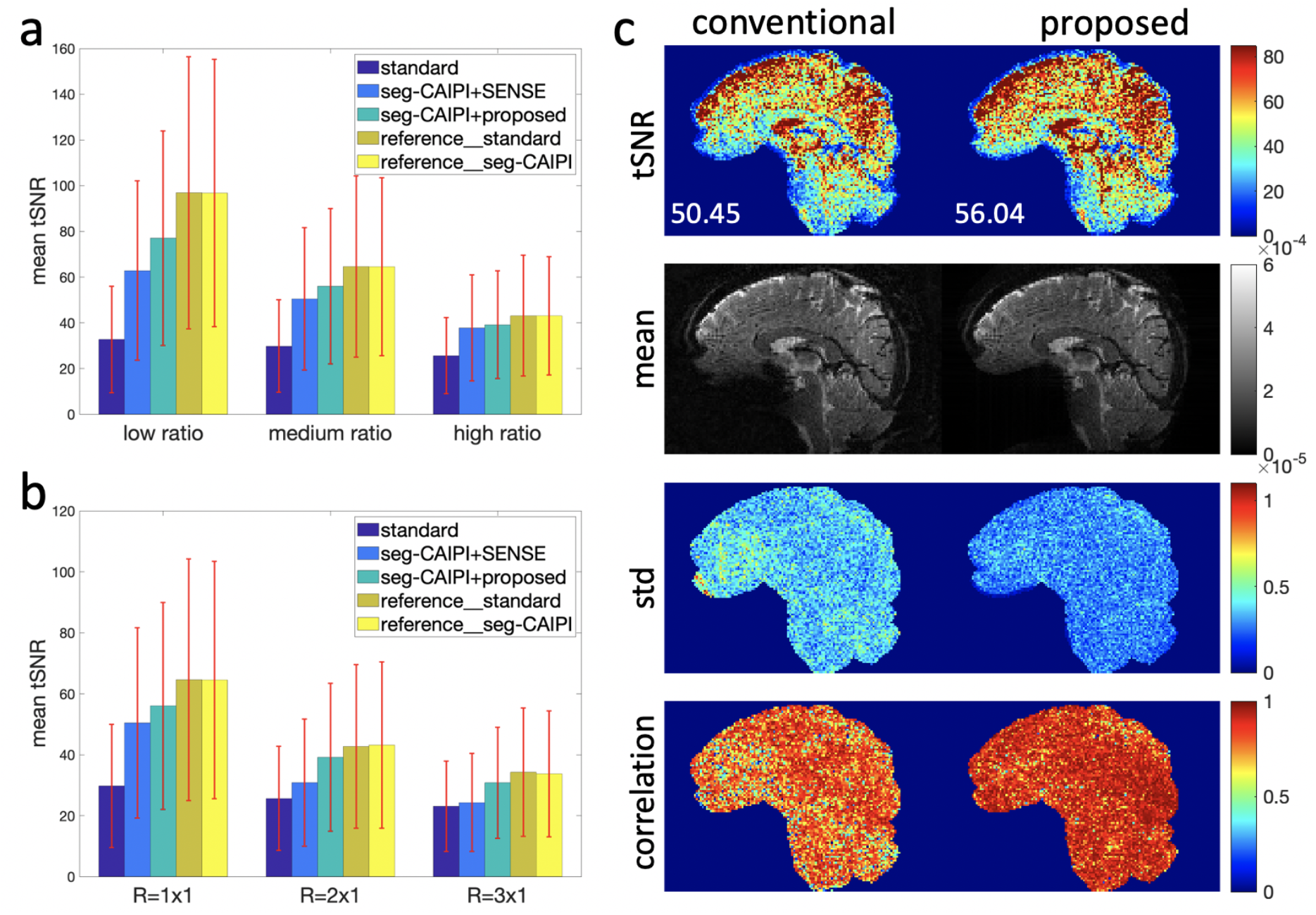
The reconstruction results of the simulation data. The dataset without inter-shot physiological variations is used as a reference. (a) Mean tSNR of different methods at varying thermal to physiological noise ratios. (b) Mean tSNR of different methods at varying acceleration factors, at medium thermal noise levels. (c) Maps of tSNR (mean value shown on the bottom left), temporal mean, standard deviation and correlation with the reference dataset of the conventional reconstruction (left) and the proposed reconstruction (right) on the seg-CAIPI data at medium thermal noise level.

Figure 4 shows the reconstruction results of a 1.2mm isotropic resolution acquisition with *R_PE_* = 3 and *R*_3*D*_ = 2 in a central sagittal slice. The mean tSNR across the whole brain (±standard deviation) was 14.81±7.70 for the standard data, and 16.81±8.20 for the seg-CAIPI data with the proposed reconstruction. In this case, the standard and seg-CAIPI(8,1) trajectories show overall similar performance with the conventional SENSE reconstruction, while the proposed reconstruction on the seg-CAIPI(8,1) data achieves a 13% improvement in mean tSNR compared to the standard data. At this high resolution, tSNR values are relatively low, and the improvement over the standard acquisition and reconstruction is limited by the thermal noise dominated regime.

**Figure 4.**
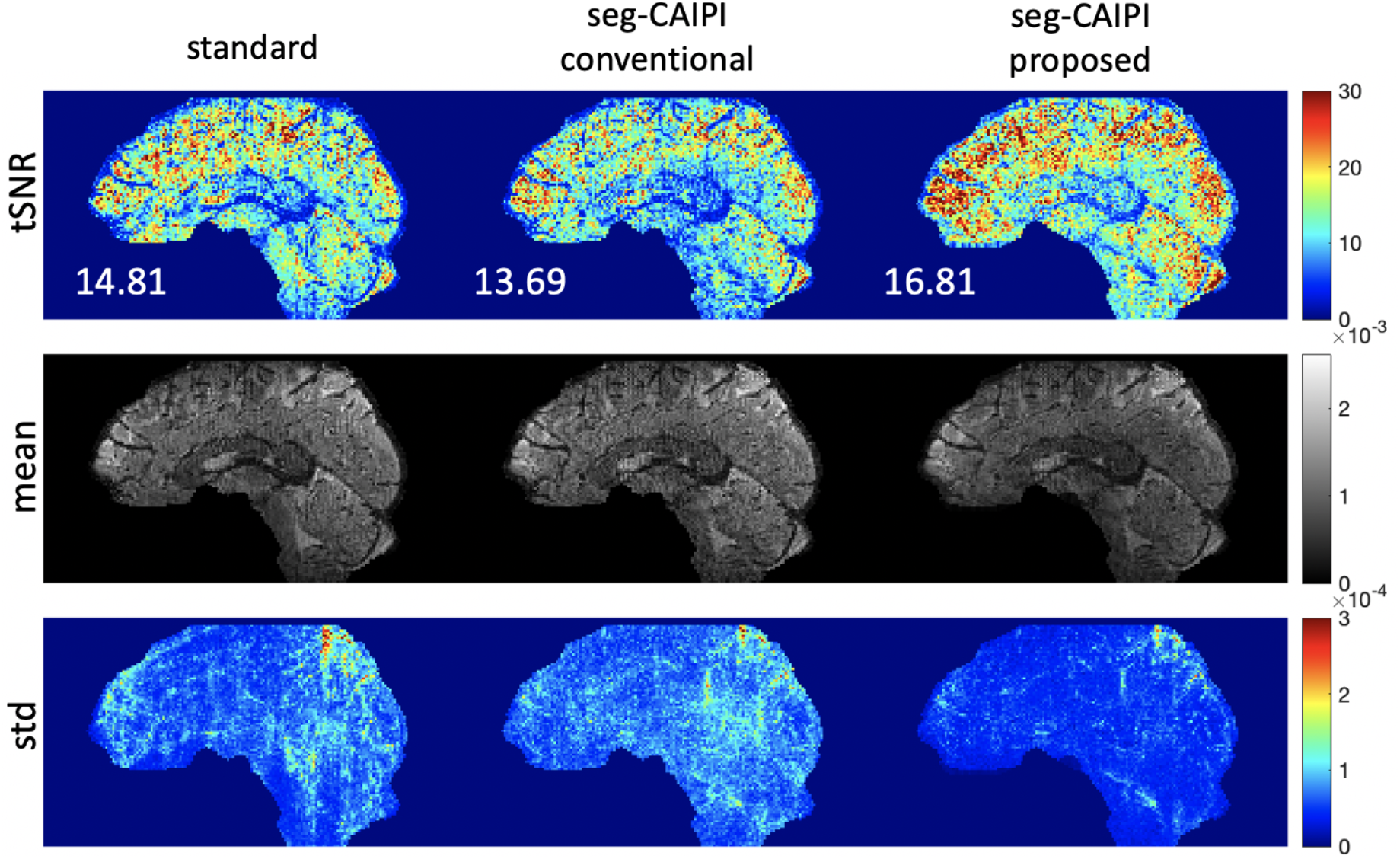
The reconstruction results of the 1.2mm isotropic resolution in-vivo dataset acquired on subject 3. The mean value across the whole brain is shown on the bottom left for each tSNR map.

Figure 5 shows the reconstruction results of a lower resolution 1.5mm isotropic resolution dataset, where we compare different seg-CAIPI trajectories: seg-CAIPI(4,1), seg-CAIPI(6,1), seg-CAIPI(8,1), along with the standard trajectory acquisition. The tSNR map and temporal mean magnitude image of the standard reconstruction are shown in Figs. 5a and 5b. The tSNR maps of different seg-CAIPI data by both conventional (left) and proposed (right) reconstructions are shown in Fig 5c. Note the first 5 image volumes of seg-CAIPI(4,1) data were excluded in tSNR calculation due to significant motion corruption. In this case, compared with the standard trajectory data, all three seg-CAIPI datasets show an improvement in mean tSNR with both the conventional and the proposed structured low-rank reconstruction. Specifically, the proposed reconstruction on the seg-CAIPI(8,1) data achieves a ~57% improvement in mean tSNR compared to the standard trajectory data with conventional reconstruction. The mean tSNR across the whole brain (±standard deviation) was 23.85±14.05 for the standard data, 37.56±19.41 for the seg-CAIPI(8,1), 34.35±18.57 for the seg-CAIPI(6,1) and 35.94 ± 19.20 for the seg-CAIPI(4,1) data with the proposed reconstruction. The corresponding temporal mean magnitude images of all the seg-CAIPI data can be found in Supplementary Figure S1.

**Figure 5.**
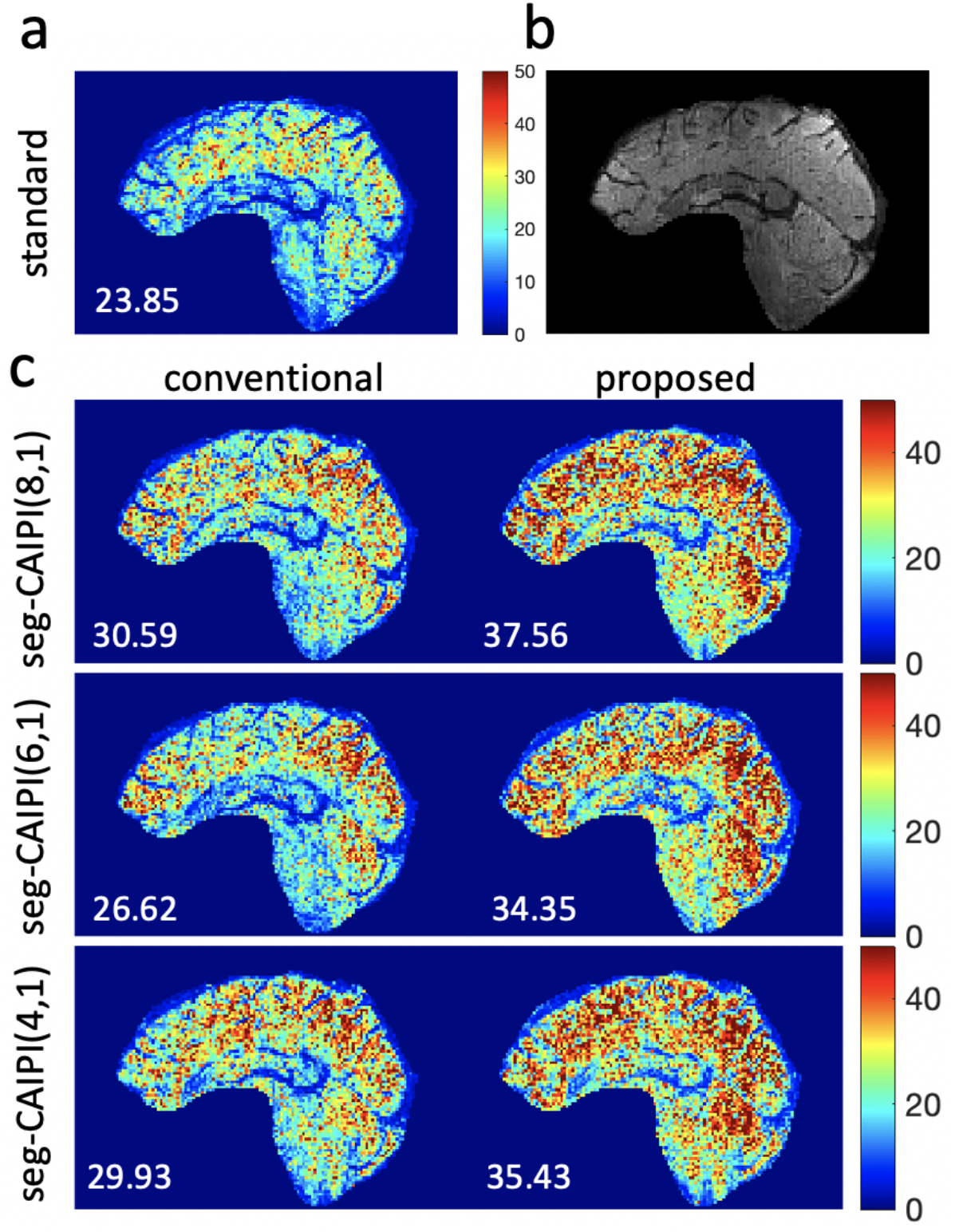
The reconstruction results of a 1.5mm isotropic resolution in-vivo dataset acquired on subject 3. (a) tSNR maps of the standard trajectory data. (b) Temporal mean magnitude image of the standard trajectory data. (c) tSNR maps of the seg-CAIPI(8,1), seg-CAIPI(6,1), seg-CAIPI(4,1) data by both conventional (left) and the proposed reconstruction (right). Mean tSNR across the whole brain is shown on the bottom left for each tSNR map.

Figure 6 shows the results of another 1.5mm isotropic resolution dataset, where we compared the seg-CAIPI data acquired with and without *k_z_* blips, i.e., seg-CAIPI(8,1) and seg-CAIPI(8,0), and the standard trajectory data. The tSNR map and temporal mean magnitude image of the standard trajectory data are shown in Figs. 6a and 6b, respectively. The tSNR maps of different seg-CAIPI data by both conventional (left) and proposed (right) reconstructions are shown in Fig 6c. In this case, the seg-CAIPI data hardly improves tSNR using conventional reconstruction, whereas the proposed reconstruction on the seg-CAIPI(8,1) data still achieves a ~65% improvement in mean tSNR compared to the standard trajectory data. However, the proposed reconstruction of the seg-CAIPI(8,0) data does show slightly reduced signal intensity, whose temporal mean magnitude image can be found in Supplementary Figure S2. The mean tSNR across the 3D whole brain (±standard deviation) is 21.26±14.30 for the standard data, 29.66±16.01 for the seg-CAIPI(8,0), 34.98±19.36 for the seg-CAIPI(8,1) data with the proposed reconstruction.

**Figure 6.**
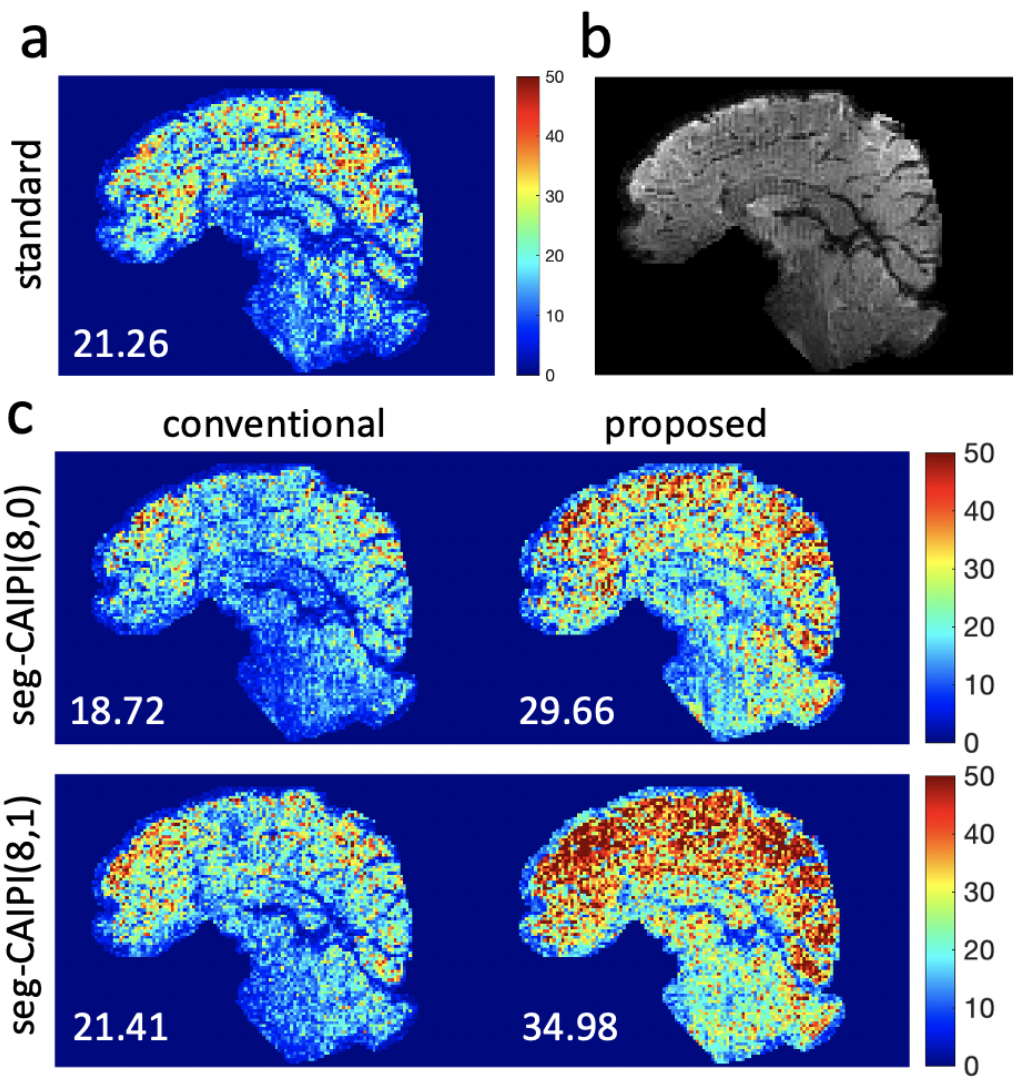
The reconstruction results of a 1.5mm isotropic resolution in-vivo dataset acquired on subject 1. a. tSNR maps of the standard trajectory data. b. Temporal mean magnitude image of the standard trajectory data. c. tSNR maps of the seg-CAIPI(8,0), seg-CAIPI(8,1) data by both conventional (left) and the proposed reconstruction (right). Mean tSNR across the whole brain is shown on the bottom left for each tSNR map.

Figure 7 shows the reconstruction results of three 1.8mm isotropic resolution datasets with undersampling factors *R*_3*D*_ = 1/2/4, and *R_PE_*= 2. In all three cases, the seg-CAIPI(8,2) data with and without the proposed reconstruction both achieve significantly higher tSNR than the standard trajectory. In particular, the proposed reconstruction on the seg-CAIPI data at *R*_3*D*_ = 4 achieves comparable tSNR to the standard acquisition at *R*_3*D*_ = 1, which suggests that the proposed method could enable an additional factor of 4x acceleration without loss of tSNR. All the corresponding temporal mean magnitude images can be found in Supplementary Figure S3. In Fig.7b, the mean power spectrum of the reconstructed time courses across the whole masked 2D slice of data acquired at R = 2 × 4 is shown, where the proposed reconstruction on the seg-CAIPI data achieved the lowest power across the entire frequency band. The peaks around 0.3Hz are likely associated with respiration. Here, the spectra are aligned to have matched energy at 0 Hz, which was realised by scaling the time course at each pixel by its temporal mean. Figure 8 shows an axial slice of another 1.8mm isotropic resolution dataset acquired using seg-CAIPI(8,2) trajectory on a different subject. Retrospective under-sampling was performed on this data to achieve an acceleration factor *R*_3*D*_ = 2 while keeping *R_PE_* = 2. Table 2 shows the mean tSNR across the whole brain (±standard deviation) of the 1.8mm isotropic resolution datasets for both subjects.

**Figure 7.**
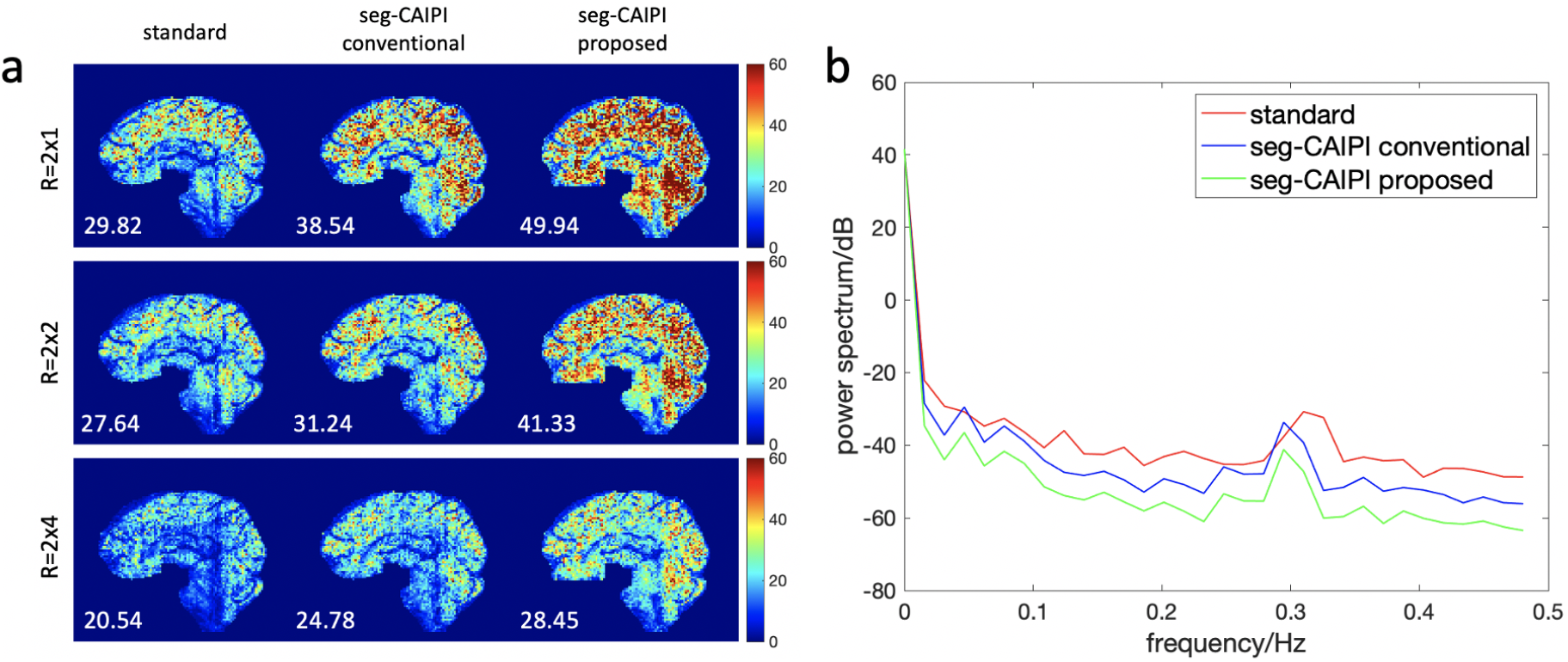
The reconstruction results of a 1.8mm isotropic resolution in-vivo dataset acquired on subject 3. (a). The tSNR maps of data acquired at R = 2 × 1/2/4 respectively. Mean tSNR across the whole brain is shown on the bottom left for each tSNR map. (b). The mean power spectrum of the reconstructed time course across the whole masked 2D slice of data acquired at R = 2 × 4.

**Figure 8.**
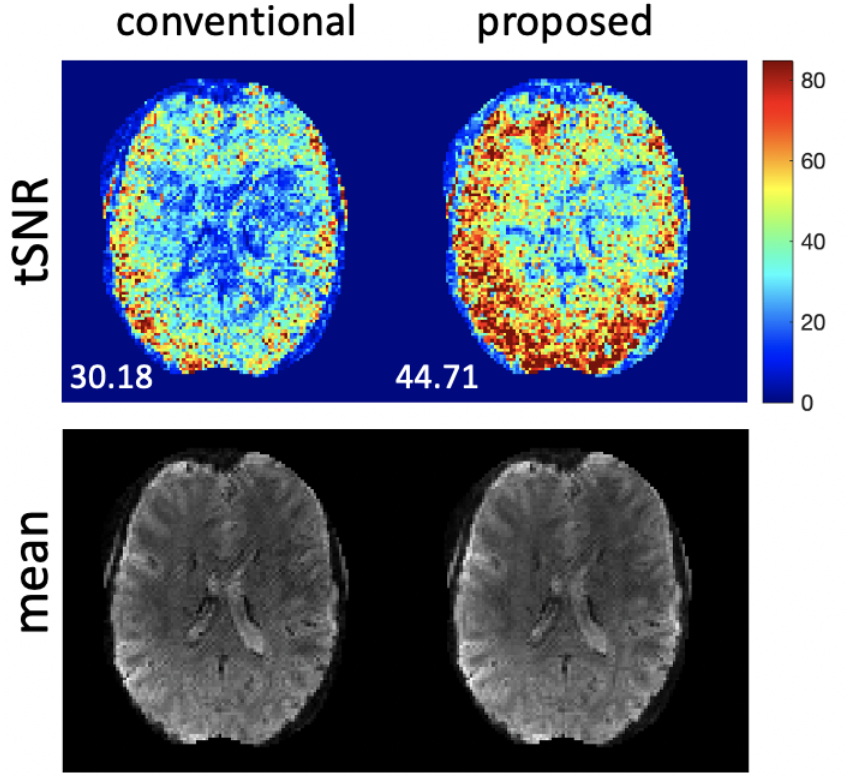
The reconstruction results of a 1.8mm in-vivo dataset at R = 2×2 (retrospective). Mean tSNR across the whole brain is shown on the bottom left for each tSNR map.

**Table 2.** The mean tSNR across the 3D whole brain (±standard deviation) of the 1.8mm isotropic resolution datasets for both subject 2 and subject 3. The dataset marked by * was retrospectively under-sampled and other datasets were prospectively under-sampled. Temporally coherent retrospective under-sampling for an R_3D_ = 2 on the standard data could completely exclude the positive/negative k_z_ data and lead to a poorly conditioned reconstruction, so it is not included for comparison here.

|  |  | standard | seg-CAIPI<br>conventional | seg-CAIPI<br>proposed |
| --- | --- | --- | --- | --- |
| subject 2 | R=2x1 | 28.09±17.20 | 40.46±22.78 | 60.70±31.31 |
|  | R=2x2* | - | 30.18±17.54 | 44.71±23.10 |
| subject 3 | R=2x1 | 29.82±19.19 | 38.54±22.84 | 49.94±28.75 |
|  | R=2x2 | 27.64±15.65 | 31.24±17.69 | 41.33±21.59 |

Figure 9 shows activation z-statistic maps from the visual task experiment at 1.8 mm isotropic resolution for different methods. Retrospective under-sampling is performed to achieve an *R*_3*D*_ = 2. The proposed reconstruction on the seg-CAIPI data demonstrates overall the highest sensitivity to activation, with clearer activation along the calcarine sulcus (indicated by green boxes in the sagittal view). The seg-CAIPI data with conventional reconstruction does not show improvement over the standard trajectory data.

**Figure 9.**
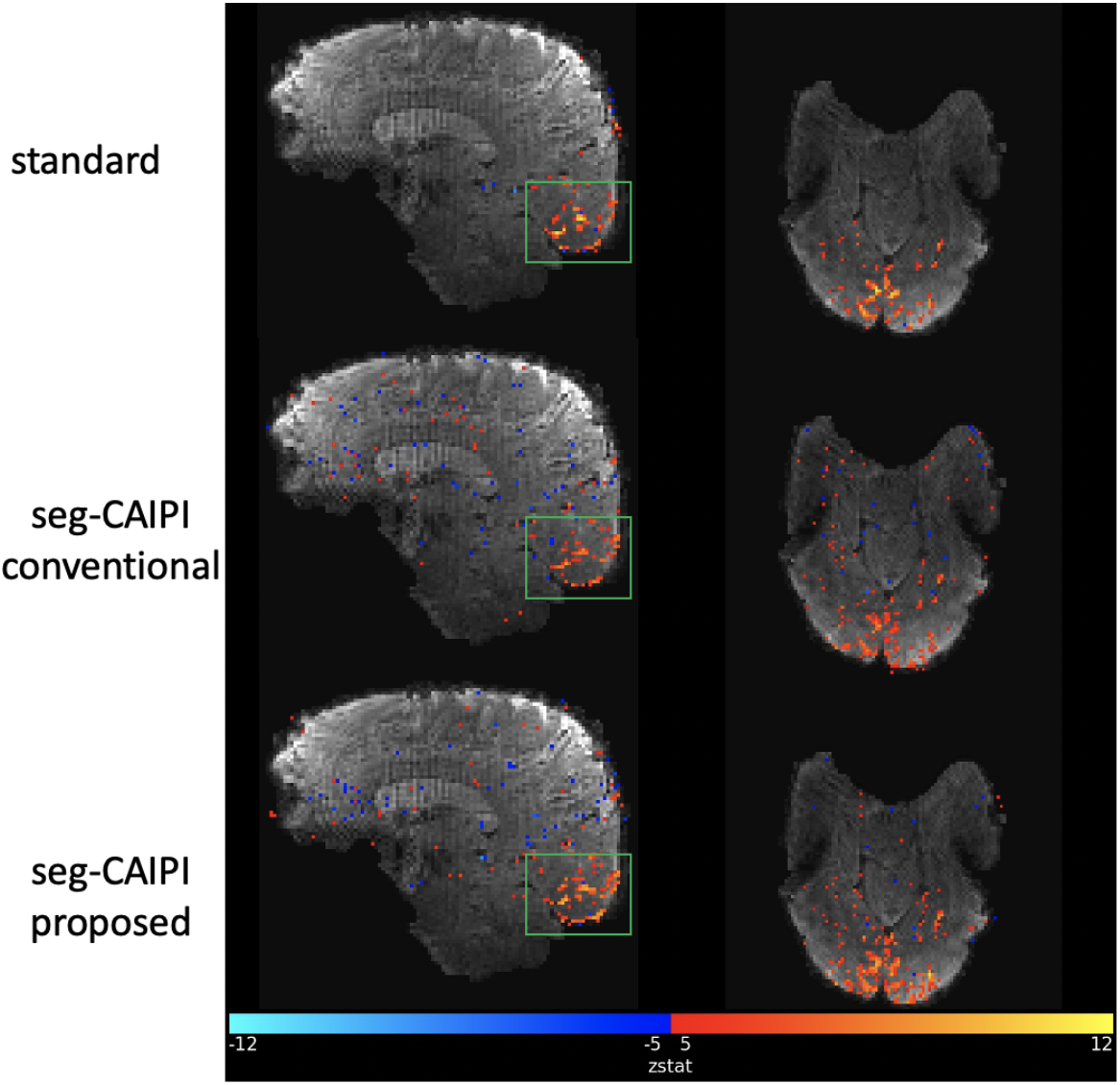
Activation maps from the flashing checkerboard experiment at 1.8 mm isotropic resolution., Activation in the calcarine sulcus is better characterised (green boxes in the sagittal view), and overall improved sensitivity to activation is observed in the proposed reconstruction on the seg-CAIPI data.

Figure 10 shows the reconstruction results of a brainstem dataset acquired at 1.2mm isotropic resolution using the seg-CAIPI(8,1) trajectory and the proposed reconstruction. The image is masked for better visualization of the brainstem. Respiration induced phase variations are expected to be stronger in brainstem than cerebrum due to the proximity to the chest cavity. An MNI template image is also shown as a reference for the anatomical structure. In both the conventional (left) and proposed reconstruction (right), fine anatomical structure in cerebellum can be observed clearly in the mean images. However, higher tSNR, particular in the cerebellum, is achieved by the proposed reconstruction.

**Figure 10.**
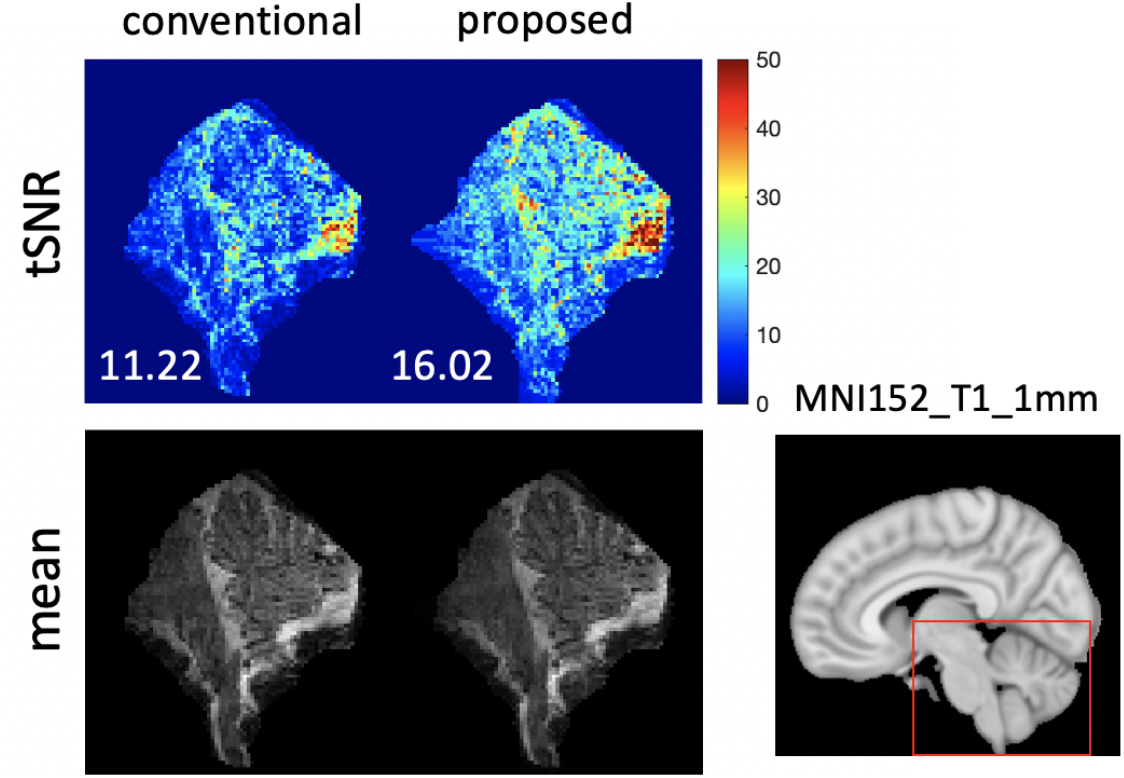
The reconstruction results of the brainstem dataset at 1.2mm isotropic resolution. Mean value across the whole masked 2D slice is shown on the bottom left for each tSNR map. An MNI152_T1_1mm template image is shown as a reference for the anatomical structure

## 4. Discussion

In this paper, we present a robust method to improve the temporal stability of 3D multi-shot EPI imaging for fMRI at 7T, which incorporates the use of a segmented 3D-CAIPI trajectory and a reconstruction method based on structured low-rank matrix completion. The seg-CAIPI sampling trajectory shows improved robustness to inter-shot phase variations compared to the standard trajectory at relatively lower spatial resolutions and acceleration factors. However, when used in conjunction with the proposed structured low-rank reconstruction, significant improvements in the resulting temporal stability compared to conventional reconstruction methods are observed in all datasets.

The seg-CAIPI trajectory is a combination of interleaved sampling and CAIPI pattern. Similar to other 3D EPI blipped-CAIPI schemes (Narsude et al., 2016; Stirnberg and Stocker, 2021; Wang et al., 2019), it acquires 3D k-space using a series of blipped, band-limited (in *k_z_*) readouts, except it returns to acquire different samples in the same “band” after each “raster” of k-space. By dividing the entire acquisition window into multiple segments, one design constraint was the optimization of the sampling trajectory for each segment, rather than for the entire sampling pattern, in order to facilitate joint reconstruction of images for each segment, instead of a single phase-corrupted image. Here, we assume intra-segment variations are negligible and consider unwanted temporal variance contributions resulting from inter-segment inconsistencies only. As the spatial distribution of the ghost artifacts is dependent on the under-sampled aliasing pattern of each segment, the spatial distribution of the unwanted temporal variance can be manipulated by altering the aliasing pattern of each segment. Thus, similar to the idea of “aliasing control” (Breuer et al., 2005) but for a different purpose, the seg-CAIPI trajectory could possibly reduce the “effective” temporal variance within the support of the object, by achieving an aliasing pattern that makes more efficient use of the FOV available. This is likely the reason why seg-CAIPI shows significantly higher tSNR than the standard trajectory, even with the same reconstruction. A simple simulation based on a numerical phantom to illustrate this can be found in the Supplementary Figure S4. Note that the efficacy of this strategy is dependent on the relative size of the object (modulated by the coil sensitivities) with respect to the FOV, and it is likely not as significant given the compact FOVs typically used. Note as the main consideration of the seg-CAIPI trajectory is the optimization of the trajectory of each segment, the overall shot-combined trajectory is not guaranteed to be optimal in terms of g-factor. However, there is no conflict between these two factors, and it is possible to further refine the seg-CAIPI trajectory by taking care of the overall shot-combined trajectory as well.

The advantage of the seg-CAIPI trajectory alone may not be robust across all experimental conditions, and are likely to be affected by factors such as spatial resolution and acceleration factors. However, when combined with the proposed reconstruction, robust improvement in tSNR were achieved. While optimal sampling for structured low-rank matrix recovery is still a topic of study (Haldar, 2014; Papadakis et al., 2015), we observed better performance for the proposed joint reconstruction of each segment with approximately uniform under-sampling for each segment, provided by the seg-CAIPI trajectory design. Note the improvement in tSNR of the proposed reconstruction is contributed mostly by the reduced temporal variance rather than a change in the signal intensity. However, slight differences between the temporal mean images of the conventional and the proposed reconstructions can be observed in some cases, which might be related to the non-optimal reconstruction parameters or sampling trajectory choice. It is also important to note that the proposed reconstruction operates on each volume independently, and only jointly reconstructs k-space segments corresponding to the same time-point. Hence, it reduces susceptibility to physiologically-induced variability, while retaining full temporal degrees-of-freedom and leaving the BOLD-related signal fluctuations and temporal characteristics intact. Furthermore, it works without requiring any information or external measurements of the physiological traces, and only requires the condition that the phase fluctuations are relatively slow with respect to the shot-to-shot TR.

For a given k-space coverage, there is a trade-off between the number of segments and the amount of data available for each segment. Increasing the number of segments can enhance the intra-segment consistency, but may make the reconstruction more challenging due to the reduced number of samples for each segment. In comparison, decreasing the number of segments might suffer from strong phase variations within the segment that can lead to phase cancellation effects. Both simulation and in-vivo experiments show that the number of segments = 6/*R*_3*D*_ or 8/*R*_3*D*_ at *R*_3*D*_ = 1 or 2 were sensible choices in this work, but this is not guaranteed to be optimal in all cases. Extending the current framework to hybrid radial-Cartesian sampling like 3D TURBINE (Chiew et al., 2016) may allow us to choose the optimal binning window for a segment retrospectively, due to the use of the golden angle sampling that provides near-uniform coverage at any window size. Radial sampling also has intrinsic robustness to temporal fluctuations, which may also benefit the robustness of 3D sampling further.

The proposed approach is based on MUSSELS (Mani et al., 2017), a method using Hankel structured low-rank matrix completion to reconstruct multi-shot diffusion weighted images. The idea of structured low-rank matrix completion has also been successfully employed in some other applications such as calibration-less parallel imaging reconstruction (Liu et al., 2021; Shin et al., 2014; Yi et al., 2021), EPI Nyquist ghost correction (Haldar and Zhuo, 2016; Lee et al., 2016b; Lobos et al., 2021), and trajectory error correction (Mani et al., 2018).The inherent linear dependency of the phase-corrupted multi-shot data has enabled us to leverage the low-rank constraint on its Hankel matrix representation, assuming image phase fluctuations driven by respiration are relatively spatially smooth, an assumption which has been employed in other work (Wallace et al., 2020). The reconstruction strategy employed here empirically estimated hyperparameters by optimizing for resulting tSNR, which provides a useful heuristic for hyperparameter tuning without requiring any additional training data or prior knowledge.

The proposed reconstruction incorporates the low-rank constraint in the SENSE-based parallel imaging formulation which uses coil sensitivities explicitly, derived from a separate multi-shot calibration dataset in this work. The moderate phase variations in fMRI typically do not lead to visible artifacts in the calculated coil sensitivities. However, in the worst case scenario when high fidelity coil sensitivity maps cannot be obtained from the multi-shot calibration data, alternative approaches can be used, such as employing the low-rank tensor representation (Hess et al., 2021; Liu et al., 2021; Yi et al., 2021) of multi-channel k-space for a calibration-less reconstruction without explicit use of coil sensitivity maps, or using a calibration consistency constraint that jointly identifies a coil-nullspace from the imaging and calibration data, without trusting either dataset completely (Lobos et al., 2021). In addition, we did not use partial Fourier sampling in this work, which is also compatible with the seg-CAIPI trajectory. When partial Fourier sampling is used, virtual conjugate k-space shots (Bilgic et al., 2019) can also be incorporated into the Hankel matrix construction. This approach was evaluated, however, our reconstruction results were not further improved compared to the results achieved without the addition of virtual conjugate shots (Figure S5). Some possible reasons may be that the baseline image phase was not sufficiently smooth, or that the reconstruction problem was sufficiently well-conditioned that an additional constraint did not significantly impact reconstruction fidelity.

We use the convex nuclear norm to enforce the low-rank property of the Hankel matrix in this paper. However, non-convex low-rank penalties have been shown to outperform convex approaches (Haldar, 2014), which was also evaluated in this study by using strict rank constraints, but our preliminary results did not show a significant difference between using convex and non-convex formulations, and for simplicity we chose to use the convex formulation for the results in this paper. Further investigations into non-convex optimizations could be an interesting direction for future research.

One limitation of the proposed reconstruction is the computation time. The reconstruction takes ~130 seconds (3.1 GHz Intel Core i7 and 16 GB RAM) per slice for a 116 × 48 sagittal matrix size and 32 sensitivity channels. However, all sagittal slices can be reconstructed independently, so the reconstruction does parallelize well with additional computational resources. The reconstruction time could probably be further reduced by code optimization, using an SVD-free rank minimization algorithm (Bilgic et al., 2019; Lee et al., 2016b), or a matrix lifting-free algorithm (Ongie and Jacob, 2017). Another limitation of this work is that no consideration of motion artifacts is taken into account. As the low-rank property of the block Hankel matrix representation of the multi-segment k-space relies on the assumption that different segments share the same image magnitude, motion induced magnitude mismatch between segments could violate this assumption. Thus, a future extension of this work could be to incorporate motion estimates in the forward model to improve the low-rankness of the block Hankel matrix representation and ultimately the robustness of 3D multishot joint reconstruction further. We anticipate that one particularly useful application of the proposed method is in brainstem and spinal cord imaging, where multi-shot 3D imaging can provide the high spatial resolution needed to resolve important structures, but where respiration induced phase variations are expected to be more significant, and motion artefacts could also be more severe.

## 5. Conclusion

In this work, we proposed to use the low-rank constrained reconstruction on the data acquired with the segmented 3D-CAIPI sampling trajectory, which is demonstrated to be less sensitive to inter-shot phase variations and thus improve the robustness of 3D multi-shot EPI for fMRI at 7T.

## Acknowledgement

The Wellcome Centre for Integrative Neuroimaging is supported by core funding from the Wellcome Trust (203139/Z/16/Z). MC and WW are supported by the Royal Academy of Engineering (RF201617\16\23, RF201819\18\92). MC also receives research support from Engineering and Physical Sciences Research Council (EP/T013133/1).

---

**Figure S1.**
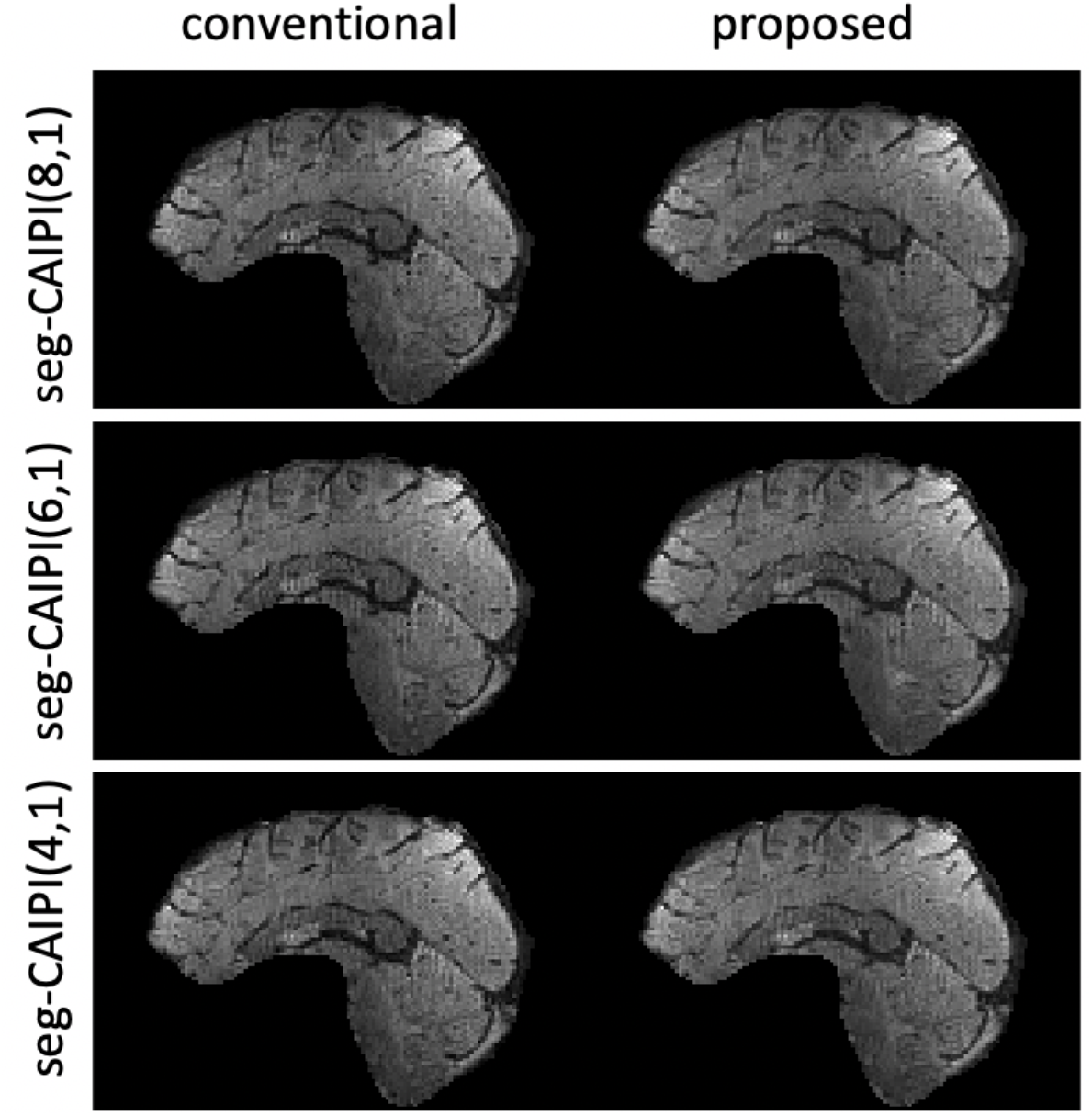
The temporal mean magnitude image corresponding to the tSNR maps of seg-CAIPI data shown in Fig.7.

**Figure S2.**
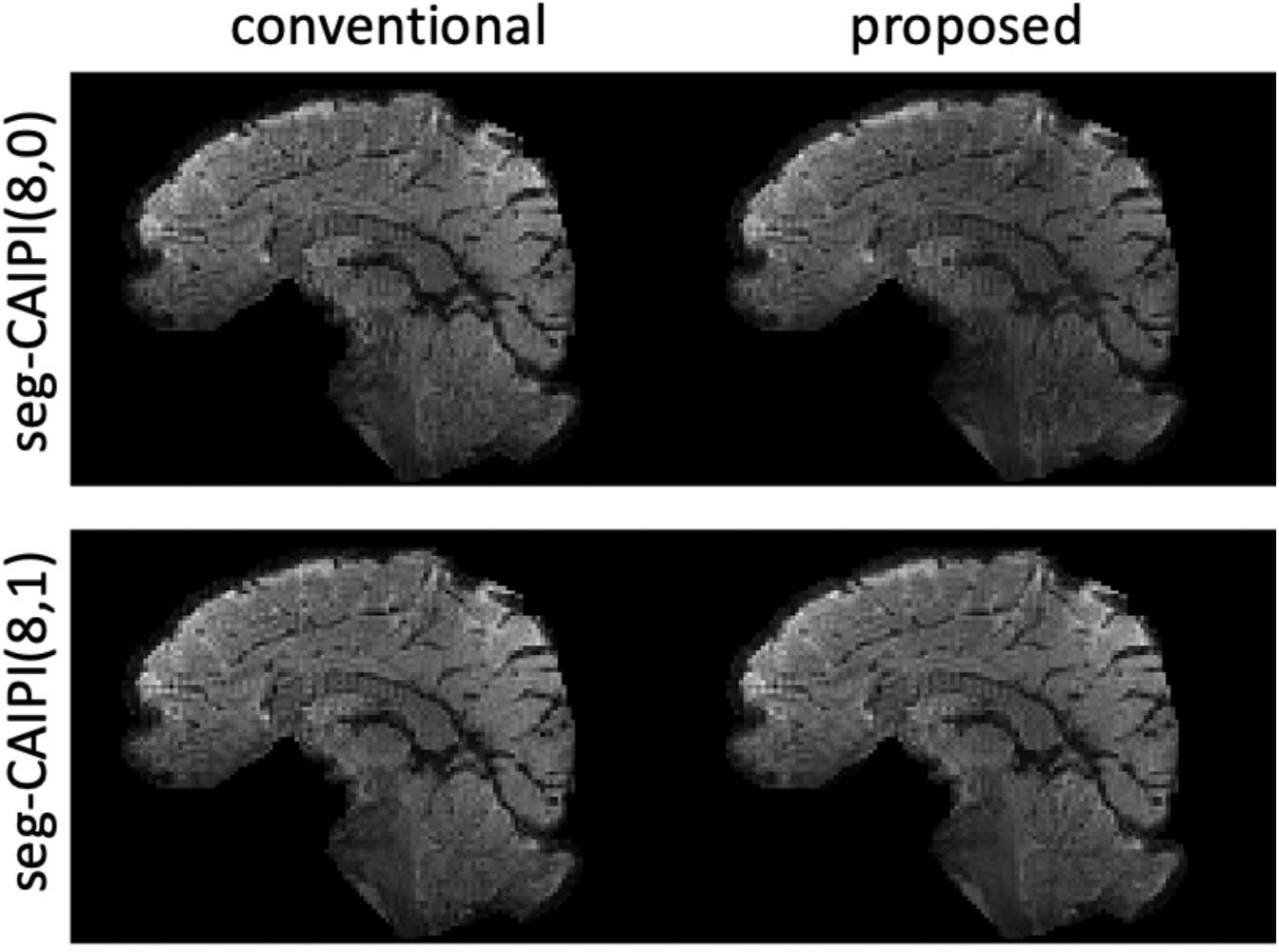
The temporal mean magnitude image corresponding to the tSNR maps of seg-CAIPI data shown in Fig.6.

**Figure S3.**
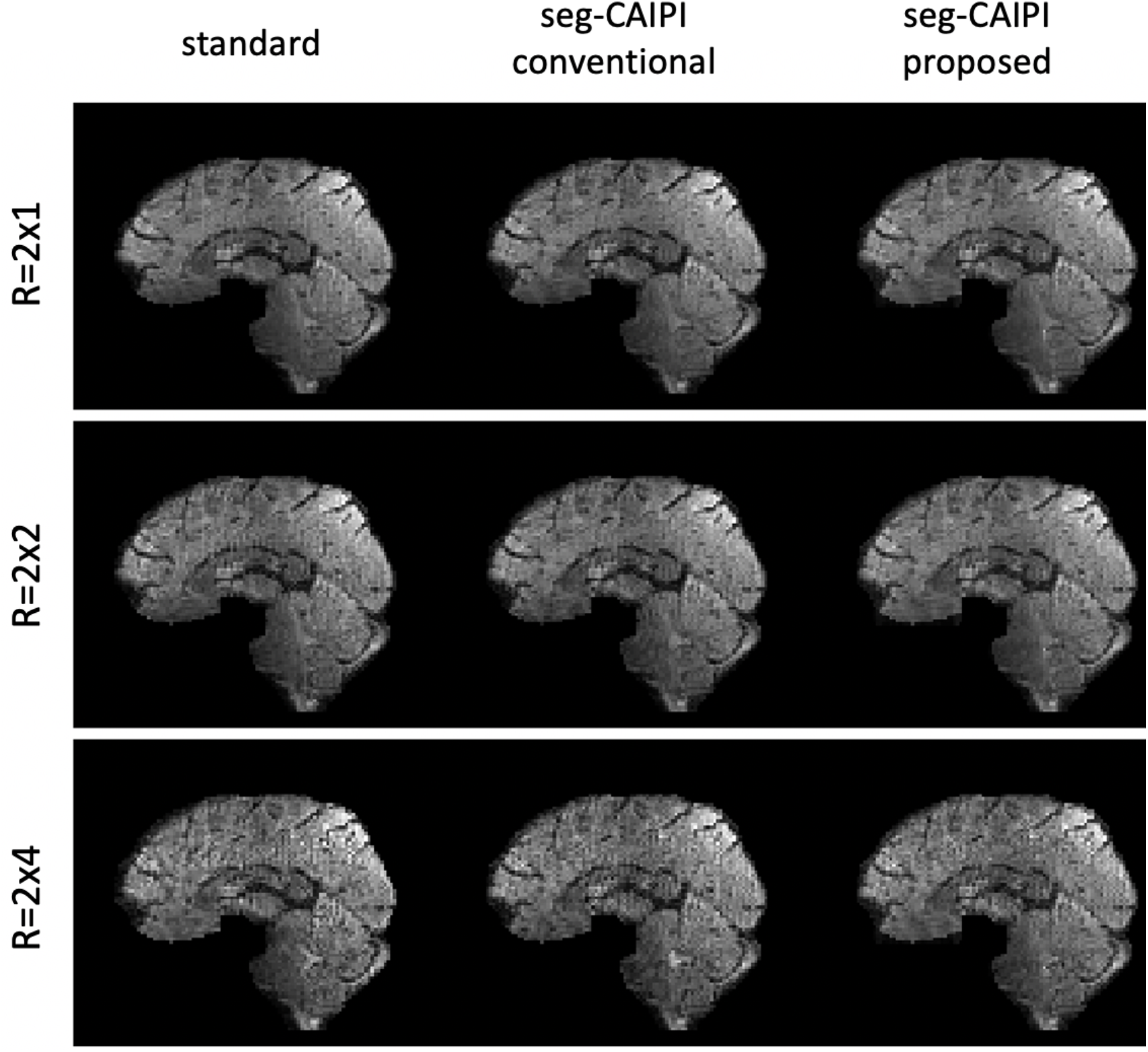
The temporal mean magnitude image corresponding to the tSNR maps shown in Fig. 7

**Figure S4.**
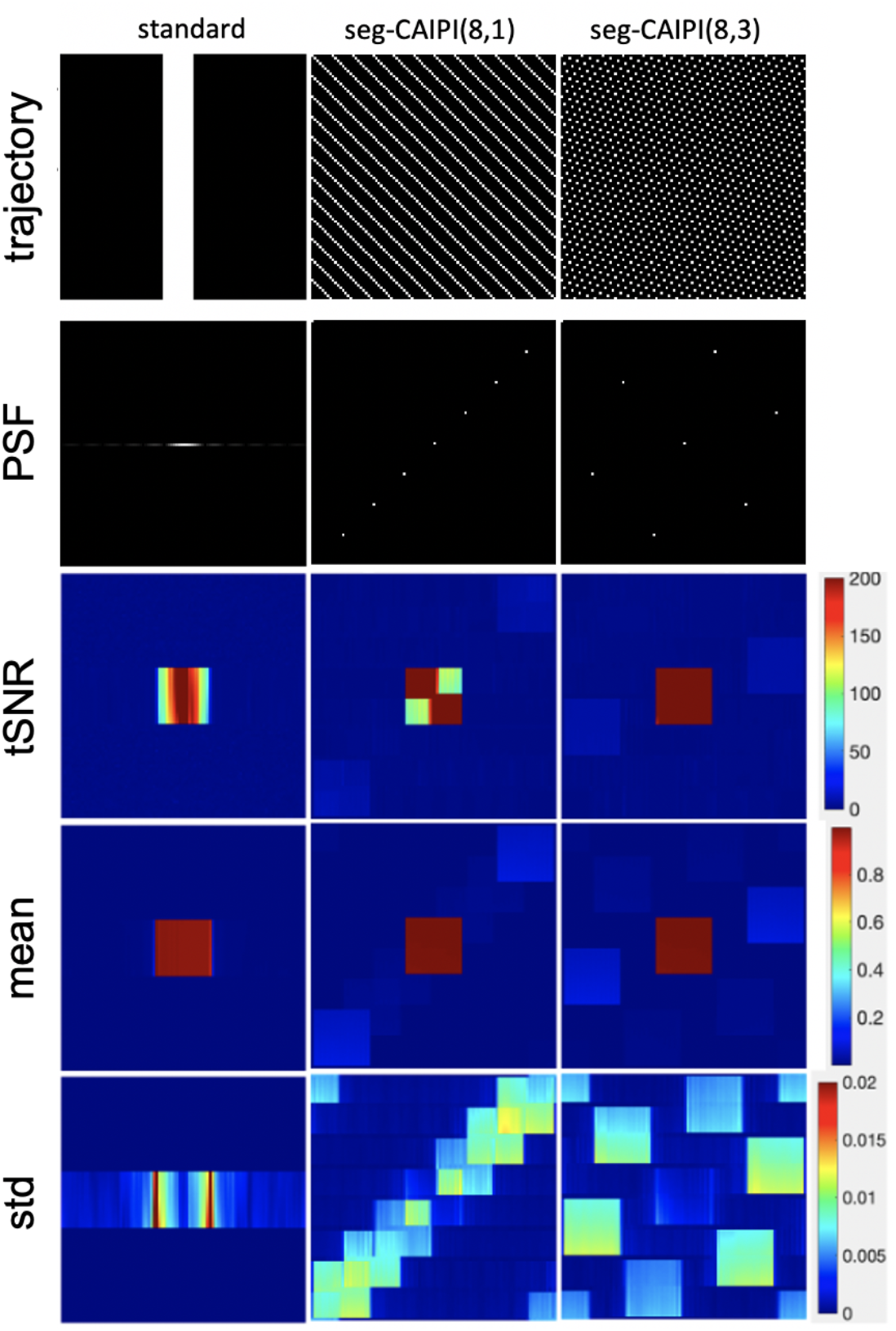
The results of a single coil numerical phantom simulation experiment. In the simulation experiment, a square numerical phantom with uniform signal intensity (see the central square in the temporal mean magnitude image) and same phase variations as in Fig.3 were used. Fully sampled multi-shot k-space datasets were generated for different trajectories. The trajectory of a segment which consists of 8 shots and its corresponding point spread function (PSF) are shown in the first and second row. The tSNR, temporal mean magnitude, temporal standard deviation maps of the reconstructed time series are shown in the third/fourth/fifth rows.

**Figure S5.**
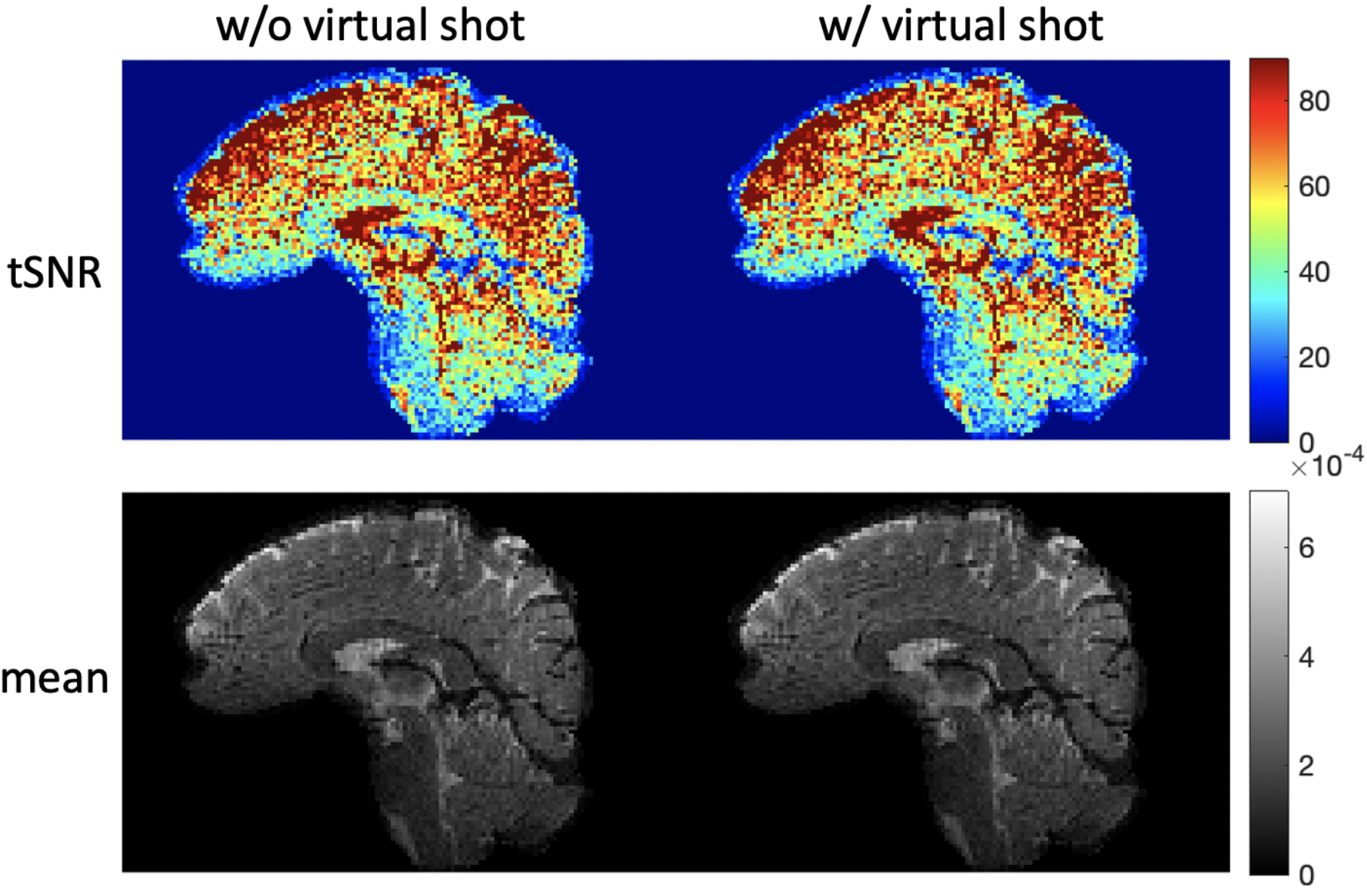
The comparison between the proposed reconstruction on the simulation data without (left, same as Fig. 3c) and with (right) using virtual shot approach. The maps of tSNR (top) and temporal mean magnitude (bottom) are compared.

